# *ETHYLENE RESPONSE FACTOR* (*ERF*) genes modulate plant root exudate composition and the attraction of plant parasitic nematodes

**DOI:** 10.1101/652321

**Authors:** Steven Dyer, Ryan Weir, Deborah Cox, Xavier Cheseto, Baldwyn Torto, Johnathan J. Dalzell

## Abstract

Plant root exudates are compositionally diverse, plastic and adaptive. Ethylene signalling influences the attraction of plant parasitic nematodes (PPNs), presumably through the modulation of root exudate composition. Understanding this pathway could lead to new sources of crop parasite resistance. Here we have used Virus-Induced Gene Silencing (VIGS) to knockdown the expression of two *ETHYLENE RESPONSE FACTOR* (*ERF*) genes, *ERF-E2* and *ERF-E3* in tomato. Root exudates are significantly more attractive to the PPNs *Meloidogyne incognita*, and *Globodera pallida* following knockdown of *ERF-E2*, which has no impact on the attraction of *Meloidogyne javanica*. Knockdown of *ERF-E3* has no impact on the attraction of *Meloidogyne* or *Globodera* spp. GC-MS analysis revealed major changes in root exudate composition relative to controls. However, these changes do not alter the attraction of rhizosphere microbes *Bacillus subtilis* or *Agrobacterium tumefaciens*. This study further supports the potential of engineering plant root exudate for parasite control, through the modulation of plant genes.

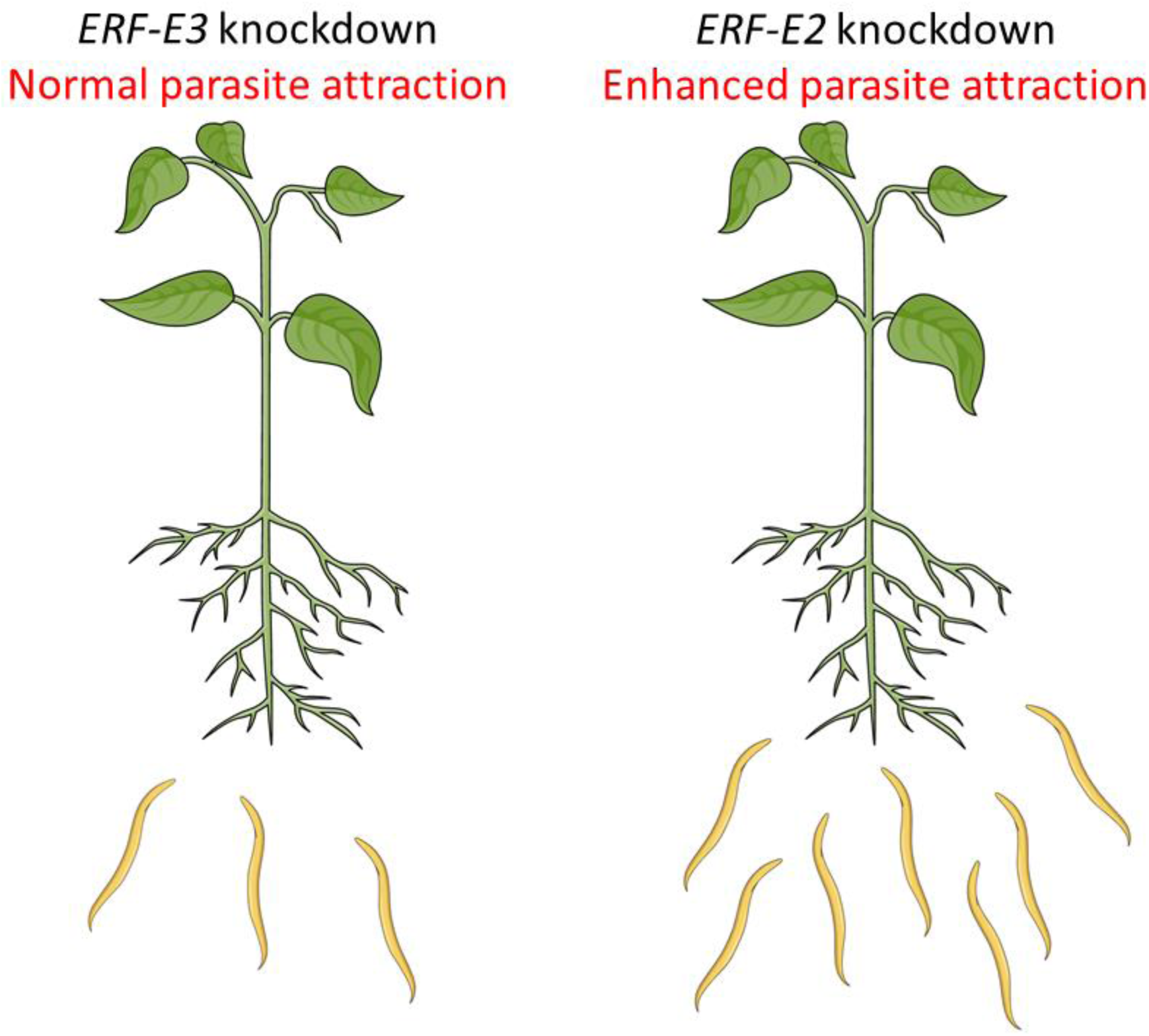

---

PPNs are estimated to reduce crop yield by 12.3% – 25% globally each year (Coyne et al., 2018; Nicol et al., 2011). The non-feeding second stage PPN juvenile (J2) hatches in the soil, and must find and invade a suitable host plant by following concentration gradients of water-soluble and volatile components of plant root exudate (Čepulytė et al., 2018; Murungi et al., 2018). We have previously demonstrated that the modulation of ABC transporter genes can alter plant root exudate composition, and parasite hatching and attraction *ex planta* (Cox et al., 2019a; Warnock et al., 2016). This represents an interesting approach to crop plant resistance, as it manipulates parasite behaviour in the rhizosphere, and limits host invasion events that can lead to secondary infection, and metabolically expensive defence responses. However, our capacity to engineer beneficial root exudate profiles is limited by a lack of fundamental insight to the complex genotype-environment-phenotype interaction that underpins this biology.

Ethylene is a gaseous plant hormone that plays a key role in regulating growth, development and physiological response to stress (Ju and Chang, 2015). Ethylene receptors are localised to the endoplasmic reticulum, and constitutively inhibit downstream signalling pathways through the action of a Ser/Thr kinase called CONSTITUTIVE TRIPLE RESPONSE 1 (CTR1). Ethylene receptor agonism interferes with receptor-mediated activation of CTR1, leading to the transcriptional activation of *ETHYLENE RESPONSE FACTOR* (*ERF*) genes. ERF proteins belong to the APETALA2/ERF transcription factor family, and regulate ethylene-responsive genes through binding to GCC-box elements (Mizoi et al., 2012). Ethylene is linked to various aspects of host-PPN interactions (Warmerdam et al., 2019; Piya et al., 2019; Wubben et al., 2004; Wubben et al., 2001), and Inhibition of ethylene signalling enhances attraction of the root knot nematode *Meloidogyne hapla*, and the cyst nematode *Heterodera glycines* to plant roots (Hu et al., 2017; Fudali et al., 2013). This indicates that constitutive receptor-mediated activation of CTR1 and negative regulation of *ERF* genes in the absence of ethylene, promotes attraction of PPNs. We therefore hypothesise that one or more *ERF* genes must regulate root exudate composition and parasite attraction. On that basis, knockdown of implicated *ERF* genes should phenocopy the elevated attraction of PPNs to root exudates, as observed when host plants are treated with ethylene synthesis inhibitors. Such genes could be exploited as novel sources of resistance through gain of function mutation, or over-expression.

*ERF-E2* and *ERF-E3* are expressed constitutively in tomato plants; however, the expression of *ERF-E2* increases during fruit development and ripening, whereas *ERF-E3* expression decreases (Liu et al., 2016). PPN species synchronise their life-cycle with that of host plants. For example, potato cyst nematodes hatch at very low frequency without the positive stimulus of host plant root exudate, and they respond differently to root exudates that have been collected from developmentally distinct host plants (Byrne et al., 2001). We further hypothesised that *ERF* genes which are activated towards the end of the host life-cycle (e.g. during fruiting - *ERF-E2*) are more likely to mediate root exudate changes that repel PPNs, which may represent an adaptive response of the PPN to ensure selection of a host with long term viability. Here we have used Virus Induced Gene Silencing (VIGS) to investigate the role of *ERF-E2* and *ERF-E3* genes in regulating root exudate composition, and PPN attraction to tomato cv. Moneymaker.

The experimental workflow and approach mirrors that of Cox et al. (2019a). 200 bp regions of *ERF-E2* and *ERF-E3* were synthesised individually, with a shared contiguous 200 bp region of the visual reporter gene, *PDS*. These DNA segments were incorporated into the binary VIGS vector, pTRV2, yielding pTRV2-*ERF-E2:PDS*, and pTRV2-*ERF-E3*:*PDS*, which were used in this study, along with pTRV2-*PDS* and pTRV1. pTRV1 and the pTRV2 variants were transformed into *Agrobacterium tumefaciens* strain LB4404, individually by electroporation. Tomato seedlings were inoculated by topical application of mixed pTRV1 and pTRV2 cultures (one of pTRV2-*ERF-E2:PDS*, pTRV2-*ERF-E3:PDS*, or pTRV2-*PDS* for each experimental mixture), following the methodology of Cox et al. (2019a). All downstream assays were conducted using root exudates collected from the timepoint with the highest level of gene knockdown, equating to week three-post inoculation for *ERF-E2*, and week four-post inoculation in the case of *ERF-E3* (Figure 1). In both cases, gene transcript level was reduced >50% relative to controls (*ERF-E2* [0.42 ±0.04, P<0.001***] and *ERF-E3* [0.39 ±0.01, P<0.001***]). Sequence alignment revealed that *ERF-E2* and *ERF-E3* shared the highest levels of non-target sequence similarity with each other, and served as off-target specificity controls for the VIGS process. Although non-target *ERF* gene expression was slightly reduced in both cases, differences were not statistically significant relative to controls (Figure 1).

**Figure 1.**
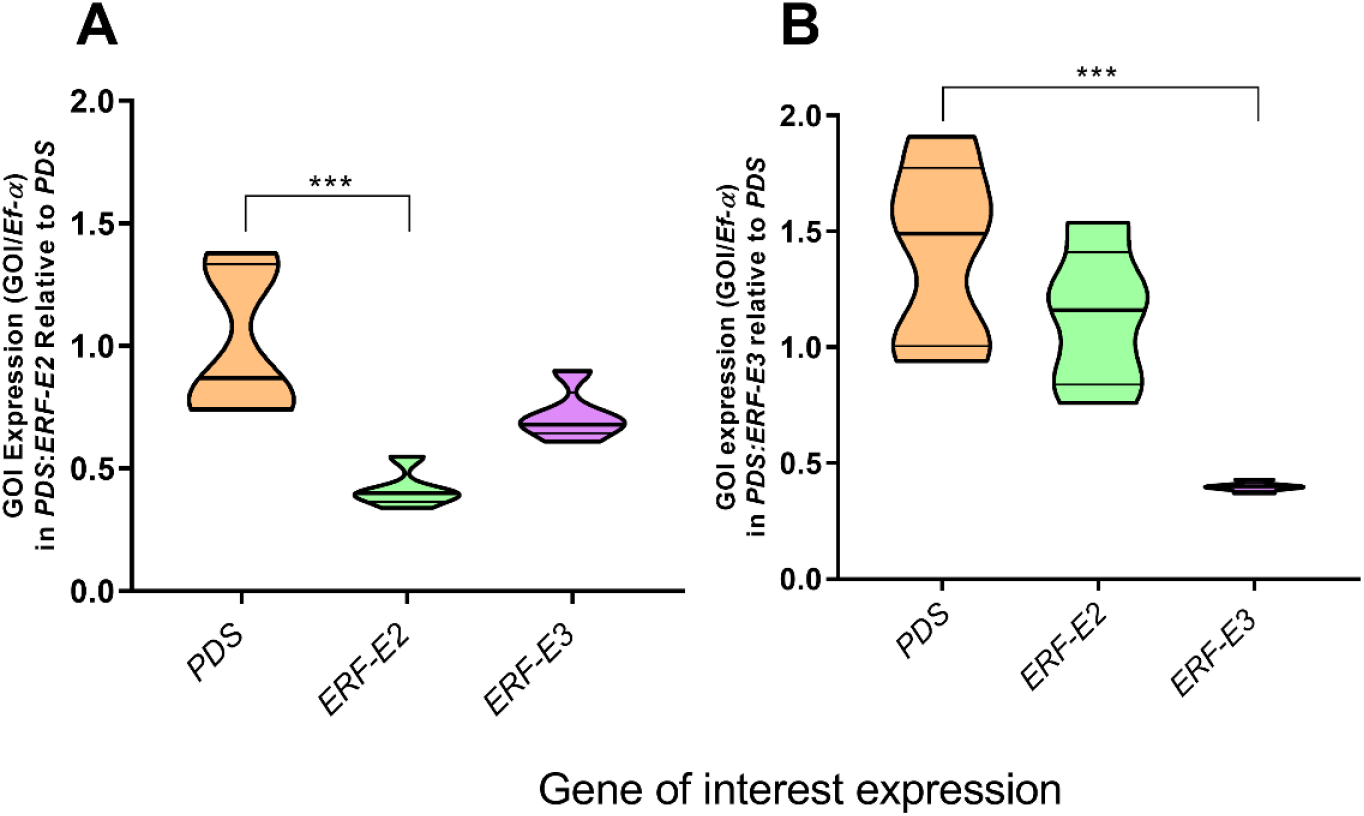
VIGS reduces the expression level of target *ERF* genes. Violin plots indicate median (inner emboldened line) and quartile (outer lines) expression values for control (*PDS* only) and experimental groups (*ERF-E2* and *ERF-E3*). (A) Target *ERF-E2* expression is significantly reduced relative to control, whereas the most similar non-target gene, *ERF-E3* was not significantly reduced. (B) Target *ERF-E3* expression is significantly reduced relative to control, whereas the most similar non-target gene, *ERF-E2* was not significantly reduced. Three VIGS-responsive plants (exhibiting leaf photobleaching following *PDS* knockdown) were pooled to produce one biological replicate, with six biological replicates used per target gene. Root tissue was gently washed free of soil, and snap frozen in liquid nitrogen. Frozen tissue was then homogenised in a pestle and mortar. Total RNA was extracted using the Maxwell® 16 LEV Plant RNA Kit and automated Maxwell® 16 AS2000 Instrument. DNA was removed using the Turbo DNase free kit (Life Technologies) according to the manufacturer’s instructions. RNA was quantified by measuring the absorbance at 260 nm. All samples were normalised to 1000 ng RNA using nuclease free H_2_O. cDNA was synthesised using the High-capacity RNA-to-cDNA kit (Applied Biosciences, UK) and diluted 5-fold prior to amplification with nuclease free H_2_O. Transcript abundance was assessed two, three, four and six weeks post-inoculation by RT-qPCR. qRT-PCR was performed in triplicate for each cDNA sample using a Rotorgene Q thermal cycler. Each individual reaction comprised 6 µl SensiFAST™ SYBR® Taq (Bioline), 0.8 µl of forward and reverse primers (*EF-α* [Solyc06g005060] F: TACTGGTGGTTTTGAAGCTG, R: AACTTCCTTCACGATTTCATCATA; *PDS* [Solyc03g123760] F: GAAGGCGCTGTCTTATCAGG, R: GCTTGCTTCCGACAACTTCT; *ERF-E2* [Solyc06g063070] F: GAAGTTCTTGCAGATCCCATATC, R: GTACATCATCGAAGGACCAAAG; *ERF-E3* [Solyc03g123500] F: GGCAAGAAGGCTAAGGTAAAC, R: CTCCACGCTGTTCATGATTG), 2.4 µl nuclease free H_2_O and 2 µl cDNA. The conditions for the reactions were as follows: 95°C × 10 min, 45 × (95°C ×10 s, 60°C × 20 s, 72°C × 25°C), 67°C->95°C rising in 1°C increments. PCR efficiencies for all reactions was calculated using Rotorgene Q software. Relative quantification of target transcript was calculated using the ^ΔΔ^Ct method relative to the endogenous housekeeping gene, *ELONGATION FACTOR 1 ALPHA* (*EF-α*). Knockdown of target genes was calculated as a ratio of expression relative to *PDS* knockdown (using pTRV2-*PDS*) in control plants. All data were assessed by one-way ANOVA and Tukey’s multiple comparison test using Graphpad Prism 8; P<0.001***.

Root exudate was collected from plants following VIGS, and was used to assess the impact of gene knockdown on parasite attraction (Figure 1 and Figure 2A). Despite *M. incognita* and *M. javanica* being closely related, our data indicate a striking variation of response to root exudates collected following *ERF-E2* knockdown; *M. incognita* displayed enhanced attraction relative to controls, whereas *M. javanica* did not. We have observed similar variation between *Meloidogyne* spp. in response to exudates collected after ABC transport gene knockdown (Cox et al., 2019a). For this line of research to develop robust and durable new sources of PPN resistance, it will be important to establish the behavioural impact that such interventions have on geographically coincident species, as well as different populations of a given species (Cox et al., 2019b). Nonetheless, our data indicate that *ERF-E2* knockdown phenocopies the enhanced attraction of certain PPN species, which is also observed following the inhibition of ethylene signalling (Hu et al., 2017; Fudali et al., 2013). It is possible that other *ERF* genes contribute to this interaction (positively or negatively), which necessitates further investigation.

**Figure 2.**
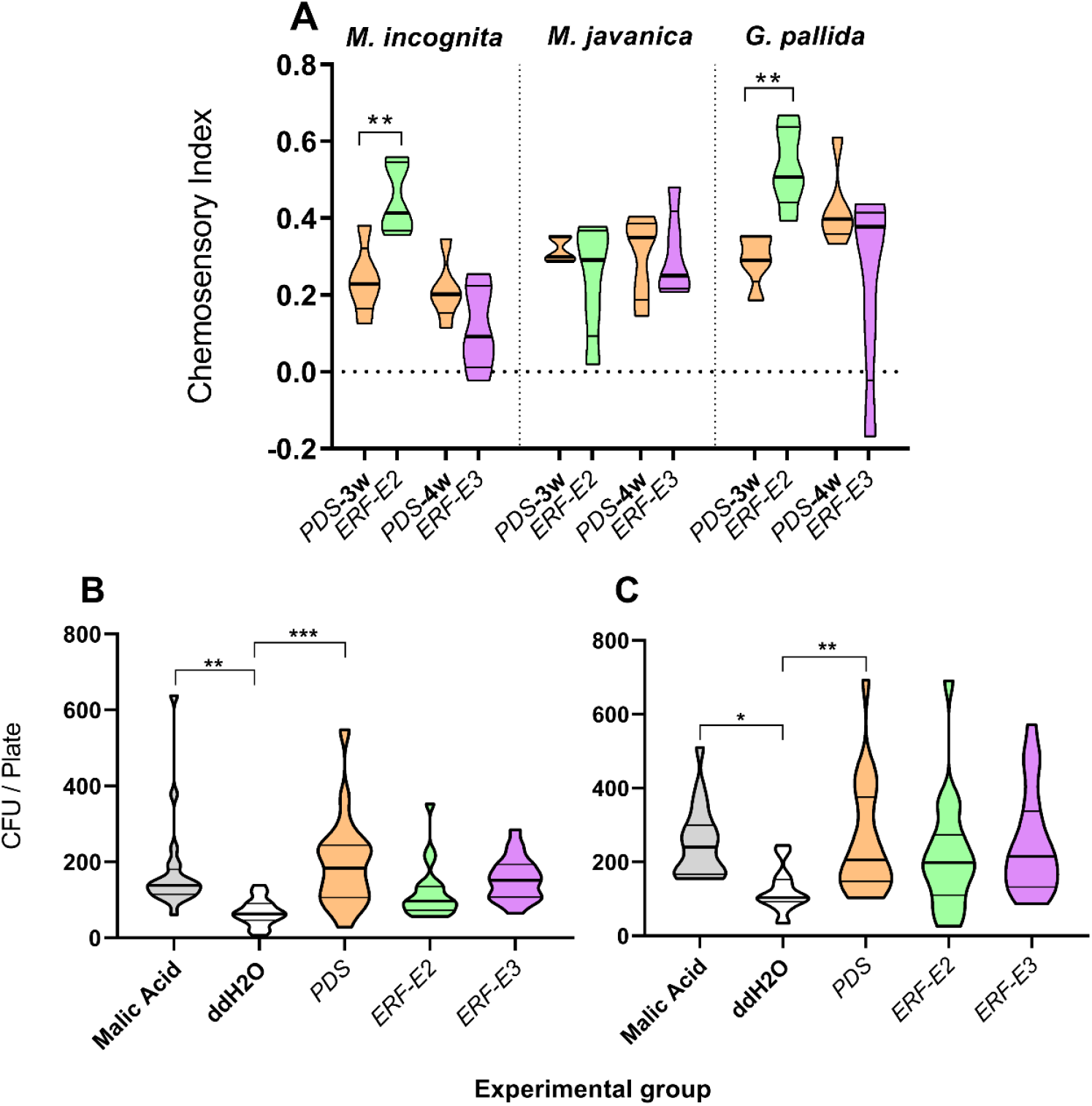
*ERF* gene knockdown modifies parasite attraction to tomato root exudate. Violin plots indicate median (inner emboldened line) and quartile (outer lines) values. (A) Knockdown of *ERF-E2* enhances the attraction of *M. incognita* and *G. pallida*, but not *M. javanica*. PPN maintenance, handling and experimentation was conducted as in Cox et al. (2019a). Briefly, *Meloidogyne* spp. were cultured in tomato cv. Moneymaker, which were maintained in a Panasonic MLR-352 plant growth cabinet (16 h light, 8 h dark, 23°C, PPFD 50 µmol·m-2·s^-1^). Eggs were hatched in sterile spring water, and used immediately for chemosensory assays. *G. pallida* was maintained on potato cv. Kerr’s Pink under field conditions in Belfast, Northern Ireland. Cysts were incubated in tomato root exudate diluted 1:1 with ddH_2_O, and freshly hatched J2s were taken immediately for chemosensory assays. Root exudates were collected from plants following gene knockdown, by submerging the root tips in 10 ml ddH_2_O for 24 h in growth chambers. The plants were removed, and the ddH_2_O / root exudate solution was passed through a 0.22 µm filter by syringe and stored at 4 °C until use. All chemosensory assays were conducted in 60 mm Petri dishes with a 3 ml base of solidified 1.5% water agar and a surface layer of 3 ml 0.5% agar slurry, as in Cox et al. (2019a). Autoclaved Whatman filter paper No. 1 discs (5 mm diameter) were saturated in 20 µl of sterile root exudate solutions, or ddH_2_O, and were placed at each end of the Petri dish immediately prior to pipetting 150 freshly hatched PPN J2s in the centre. The assays were maintained at room temperature in the dark to allow the nematodes to migrate. After 16 hours the number of J2s residing in each zone of the assay arena were counted and used to calculate a chemosensory index, as a measure of net attraction / repulsion from the positive root exudate disc. Those in the central dead zone were not counted. (B) Knockdown of *ERF* genes has no statistically significant impact on the attraction of *A. tumefaciens* or (C) *B. subtilis* to experimental root exudates. Microbial chemotaxis assays were conducted as in Cox et al. (2019a) using *B. subtilis* (168) and *A. tumefaciens* (AGL-1), relative to ddH_2_O (negative control), malic acid (positive control, and week 3 *PDS* exudate (VIGS control). 1 µl microcapillary tubes (Sigma-Alrdich) were filled with either experimental root exudates (*ERF-E2, ERF-E3*, or *PDS*), positive (1 mM malic acid), or negative (ddH_2_O) controls. The microcapillary tubes were then placed into a microbial cell suspension (within a 96-well plate) for 1 h, and maintained at 23°C. Following the assay timecourse, the capillary tubes were removed and excess cell suspension was removed from the outside of each capillary tube by rinsing briefly with ddH_2_O. The 1 µl content of each capillary tube was ejected into 99 µl of chemotaxis buffer by positive pressure. 20 µl of each solution was spread onto a 1.5% LB agar plate. LB plates were sealed with parafilm, and incubated at 28°C for *A. tumefaciens*, or 37°C for *B. subtilis* for 48 h. Colony forming units (CFUs) were counted for each replicate plate. All data were assessed by one-way ANOVA and Tukey’s multiple comparison test using Graphpad Prism 8. P<0.05*, P<0.01**, P<0.001***.

We also sought to investigate the impact that *ERF* gene knockdown had on the attraction of economically relevant rhizosphere microbes to collected root exudates (Figure 1B, and 1C). Active chemotaxis is important for initial colonisation of plant roots by the Plant Growth-Promoting Rhizobacterium (PGPR) *B. subtilis* (Allard-Massicotte et al., 2016), and the crown gall pathogen, *A. tumefaciens* (Merritt et al., 2007). Our data indicate that the attraction of *B. subtilis* and *A. tumefaciens* is not altered relative to *PDS* controls. This suggests that it may be possible to develop novel sources of crop parasite resistance, through the manipulation of root exudate composition, without affecting other important plant-rhizosphere interactions.

The GC-MS dataset indicates significant compositional changes in root exudates following knockdown of *ERF-E2* or *ERF-E3* (in addition to *PDS*), relative to control root exudates (*PDS* knockdown only). However, it is important to note that our analysis does not encompass the full spectrum of root exudate chemistry. Although we have demonstrated that octadecanoic acid is an inhibitor of PPN attraction in previous work (Cox et al., 2019a), our data suggest that it is not a biologically relevant inhibitor following *ERF-E2* knockdown, despite being significantly elevated, along with 2,3-dimethylpropyl octadecanoate (Figure 3). This suggests that other compositional changes have a greater influence on PPN behaviour in this context; similar observations have been documented previously (Kihika et al., 2017). The GC-MS data also demonstrate a substantial developmental influence on exudate composition, with control exudates exhibiting clear differences between week 3 and week 4-post inoculation. This agrees with previous work on the link between developmental stage and root exudate composition (Chaparro et al., 2013). Collectively, our data suggest that *ERF* gene knockdown may have widespread impacts on transcriptional networks and metabolic pathways contributing to exudate composition. Further work will be required to identify genes that are regulated by ERF-E2 (and other ERF proteins), which would allow more specific and targeted interventions for exudate modification and parasite resistance.

**Figure 3.**
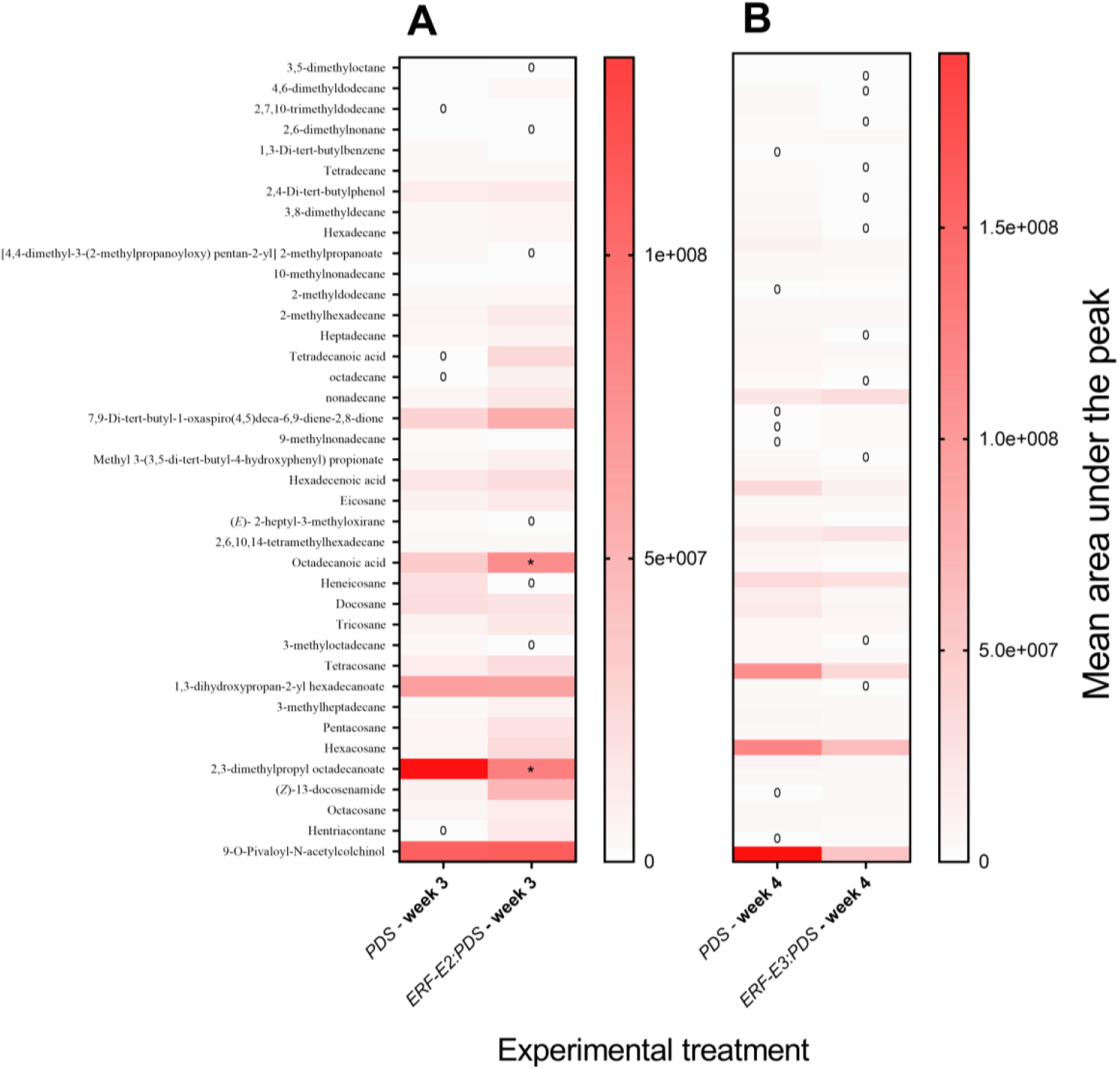
Heatmap showing differences in the relative abundance of identified root exudate compounds across experimental exudates. The mean composition of 10 biological replicates (three plants per replicate) is plotted for each experimental group post-VIGS, and has been assessed by two-way ANOVA, and Tukey’s multiple comparison test. ‘0’ indicates that the compound was not detected. Statistical significance is indicated relative to the *PDS* knockdown control, P<0.05*. Root exudates were freeze dried and stored at -80°C until processing (10 biological replicates for each treatment group). Samples were extracted with GC-grade dichloromethane (1 mL) (Sigma–Aldrich, USA), vortexed for 10 s, sonicated for 5 min, and centrifuged at 14,000 rpm for 5 min. The organic phase was dried over anhydrous Na_2_SO_4_, concentrated to 50 µL under a gentle stream of N2 and then analysed (1.0 µl) by GC-MS on a 7890A gas chromatograph linked to a 5975 C mass selective detector (Agilent Technologies, USA). The GC was fitted with a HP5 MS low bleed capillary column (30 m × 0.25 mm i.d., 0.25 µm) (J&W, Folsom, CA, USA). Helium at a flow rate of 1.25 ml min^-1^ served as the carrier gas. The oven temperature was programmed from 35 to 285°C with the initial temperature maintained for 5 min then 10°C min^-1^ to 280°C, held at this temperature for 20.4 min. The mass selective detector was maintained at ion source temperature of 230°C and a quadrupole temperature of 180°C. Electron impact (EI) mass spectra were obtained at the acceleration energy of 70 eV. Fragment ions were analysed over 40–550 m/z mass range in the full scan mode. The filament delay time was set at 3.3 min. A HP Z220 SFF intel xeon workstation equipped with ChemStation B.02.02. acquisition software was used. The mass spectrum was generated for each peak using Chemstation integrator set as follows: initial threshold = 5, initial peak width = 0.1, initial area reject = 1 and shoulder detection = on. The compounds were identified by comparison of mass spectrometric data and retention times with those of authentic standards and reference spectra published by library–MS databases: National Institute of Standards and Technology (NIST) 05, 08, and 11.

Our data indicate that *M. incognita* and *G. pallida* regulate their behaviour *ex planta* in response to *ERF-E2*, but not *ERF-E3*. This corroborates our hypothesis that one or more *ERF* gene products regulate root exudate composition, phenocopying the enhanced attraction of PPNs observed following the inhibition of ethylene synthesis (Hu et al., 2017; Fudali et al., 2013). Our data also demonstrate that *M. incognita* and *G. pallida* use developmentally regulated signalling processes in the host plant to coordinate host selection behaviour *ex planta. ERF-E2* expression increases during host fruiting, whereas *ERF-E3* expression decreases (Liu et al., 2016). *M. javanica* does not however respond in the same way as the other species, which warrants further investigation. Collectively, our data point to the potential application of *ERF* genes, or genes that are regulated by ERF transcription factors in the manipulation of parasite behaviour in the rhizosphere. It may be possible to enhance the efficacy of trap crops, or push-pull strategies through breeding or biotechnology that exploits this line of research. Encouragingly, we do not observe any impact on the behaviour of non-target soil microbes, suggesting that parasite-specific interventions may be possible. Considerable progress has been made in developing novel sources of resistance to the parasitic plant, *Striga* spp., through the manipulation of root exudate composition (Gobena et al., 2017; Samejima et al., 2016; Jamil et al., 2011;). Whilst there is still much unknown of how PPNs perceive and respond to plant root exudates, recent work continues to provide valuable insight to ther *ex planta* biology and behaviour (Bell et al., 2019; Cox et al., 2019a; Cox et al., 2019b; Tsai et al., 2019; Čepulytė et al., 2018; Hoysted et al., 2018; Kirwa et al., 2018; Warnock et al., 2016).

